# Genetically based variation in fitness and carbon assimilation among bur oak populations

**DOI:** 10.1101/2024.10.30.620350

**Authors:** Lucy M.S. Rea, Laura Ostrowsky, Rebekah Mohn, Mira Garner, Lindsey Worcester, Cathleen Lapadat, Heather R. McCarthy, Andrew L. Hipp, Jeannine Cavender Bares

## Abstract

Ongoing climate change will negatively impact tree populations unless they are able to acclimate to the changes in their local environment. Effective planning for climate adaptation management requires an understanding of the current state of local adaptation and physiological performance to assess whether populations are at risk of local extinction, to determine if seed movement is appropriate, and to select appropriate seed sources if intervention is needed. We established a new reciprocal transplant experiment (ACE, Adaptation to Climate and Environment) across a latitudinal gradient in North America to examine the impacts of warming on three bur oak (*Quercus macrocarpa*) populations across much of the species’ range. We established common gardens in Minnesota, Illinois, and Oklahoma with seedlings grown from seeds collected within 50 km of each of those locations from a total of sixty maternal families. We aimed to 1) assess local adaptation in each of the populations using survival and size as fitness metrics, and 2) evaluate physiological responses to different environments along the latitudinal gradient. We found that northern populations are maladapted to hotter climates as evidenced by their low survival, growth, and photosynthetic rates in the warmest common garden. The southernmost population had the highest survival rate, growth rate, and fitness of the three populations in the southernmost garden, providing evidence for local adaptation to the warmest site. However, conditions in the middle garden resulted in the highest fitness and best physiological performance for all populations. Growth and survival were correlated in the middle garden but were decoupled in the northern and southern gardens. This decoupling is likely due to stress associated with more extreme climates at the ends of the gradient that led to greater resource allocation to survival than to growth. Our results suggest that southern seed sources may perform well in warmer conditions in the north brought on by climate change, which has important implications for managers assisting broadly ranged tree species in adapting to climate change.

## Introduction

Warming temperatures associated with climate change will have implications for the long-term persistence of locally adapted populations as they may experience adaptational lags that cause them to become maladapted to the novel climate conditions, and may perform better at higher latitudes outside their historic range (Browne et al., 2019; Etterson et al., 2020; Gorton et al., 2022; Sáenz-Romero et al., 2017). Populations that have higher fitness in their local environment than foreign populations or higher fitness in their home environment than away environments are said to be locally adapted (Kawecki & Ebert, 2004; Williams, 1966). Local adaptation in large populations of widespread species can help maintain genetic variation within the species, which is critical in species’ persistence through changing climatic conditions, and understanding local adaptation can provide useful information that can help improve adaptative potential of populations not currently at high risk of extinction (Meek et al., 2023; Whitlock, 2015).

Trees face major threats from rapid climate change and may be particularly at risk due to their long generation time that limits their capacity to adapt quickly (Aitken et al., 2008). Given that forests are estimated to be the largest terrestrial carbon sink on the planet, reforestation is one of the most effective carbon drawdown solutions proposed, and it is especially critical to avoid losses of tree species and their genetic diversity, which provides the basis for adaptation to novel environments (Joshi et al. 2001,Bastin et al., 2019; Pan et al., 2024). Models predict that habitat suitable for temperate forests will expand northward (Rehfeldt et al., 2012), however the distance that seed and pollen can move may slow the rate at which trees themselves are able to expand. Dispersal limitations on the movement of tree species will necessitate population-level responses to novel climates through phenotypic plasticity or adaptation (Franks et al., 2014; Hoffmann & Sgrò, 2011; Huntley, 1989). Adaptational lags coupled with dispersal limitations may necessitate anthropogenic intervention to ensure survival of populations that cannot tolerate novel climate regimes in their current environment. Often, locally adapted genotypes are selected for planting in reforestation and regeneration efforts, however shifting climates may undercut the common assumption of “local is best” (Bower et al., 2024; Joshi et al., 2001; Pike et al., 2020).

Seed transfer (also referred to as population assisted migration), the practice of moving populations within their existing species range, may support locally adapted populations that have become maladapted to their current environment due to climate change. Many foresters anticipate the need for plantings in the near future that improve their forests’ ability to persist through climate change impacts (Clark et al., 2024). Movement of seeds can increase genetic diversity and the probability of survival in changing environments. However, data is needed to understand which populations and genotypes may be maladapted to climate change, and which have adaptations suited to novel conditions brought on by climate change (Xu & Prescott, 2024).

Local adaptation can be tested with reciprocal transplant experiments, which separate genetic from environmental (plastic) influences on phenotypes (Kawecki & Ebert, 2004; Schwinning et al., 2022). We established a reciprocal transplant experiment using the deciduous, broad-leaved tree species *Quercus macrocarpa* L. across a latitudinal gradient by collecting seeds from maternal families across the gradient to establish the ACE (Adaptation to Climate and Environment) experiment. *Quercus macrocarpa* is a useful species for such an experiment because it has a broad latitudinal and ecological range. Consequently, it is likely to show adaptive differentiation across its range of contrasting climatic conditions. *Quercus macrocarpa* is also an important species in oak savannas, a rare and threatened ecosystem, due to its fire resistance capabilities (Hoekstra et al., 2005). Furthermore, *Quercus* species are ecologically important tree species supporting wildlife diversity as they are an important food source for birds, insects, and mammals (Tallamy, 2021). *Quercus* species are also relevant in a management context, as they are the most common genera planted in adaptation plantings by forest managers (Clark et al., 2024). By understanding local adaptation in populations of wide-ranging tree species, we can provide valuable information on seed sourcing for land managers in seed transfer efforts, which can be a critical strategy in helping to prevent species extinction and economic losses and to maintain ecosystem services (Abrams et al., 2021; Krech et al., 2004; Williams & Dumroese, 2013). Our goals are to understand the underlying mechanisms of local adaptation in *Quercus macrocarpa* populations by taking a novel approach to integrate ecophysiology, spectral reflectance, and fitness. We do this by measuring high-throughput ecophysiological traits and processes that provide insight into the performance of our three populations in different environments and to combine survival and growth in a single measure of fitness using aster models (Geyer et al., 2007; Shaw et al., 2008).

In this study, we tested for local adaptation and adaptive differentiation by reciprocally transplanting seedlings from three populations in field conditions. We collected acorns from individual trees (half-sib families) within three populations (Minnesota, Illinois, and Oklahoma) across a climatic gradient which spans a mean annual temperature gradient of 6 –16 ᵒC. ACE was designed to reciprocally plant seedlings from each family within each population into gardens in those three states using a quantitative genetics design, such that seedlings from all families were randomized in replicated blocks in each garden. We addressed the overarching question: Do *Q. macrocarpa* progeny from three bur oak tree populations have higher fitness and physiological performance in the garden closest to their origin, or are they better adapted to a climate that has shifted northward due to warming with climate change? If populations are locally adapted, we expect them to have higher fitness and adaptive physiological performance when they are in their home garden (home vs. away), and to have higher fitness than other populations in their home garden (local vs. foreign) (Appendix S1 Fig. S1). On the other hand, if populations are adapted to a climate that has shifted northward due to the 1.1 °C warming in the past century (IPCC, 2023), we expect them to have higher fitness and physiological performance in the more northern (higher latitude) gardens compared to the hotter and drier southern (lower latitude) gardens, but northern populations to exhibit lower fitness and higher physiological stress responses in the southern garden. Furthermore, we sought to understand mechanisms behind differences in populations’ fitness in different environments by tracking differences in physiological responses and phenological timing. We expect that the physiological responses will demonstrate adaptive differentiation by providing evidence for genetic differences among populations in functional traits that are advantageous in their home environment. We expect that genotypes and populations from the south where growing seasons are longer and cold stress is reduced will exhibit physiological traits aligned with extended phenology that confers longer leaf life spans (Chabot & Hicks, 1982; Kikuzawa, 1991; Wright et al., 2004). We also anticipate that these southern genotypes will maintain productivity through hot, arid conditions (Oram et al., 2023).

In contrast, in populations from higher latitudes with shorter growing seasons we expect genotypes to have higher photosynthetic rates and higher relative growth rates (see Appendix S1 Fig. S2). Higher carbon assimilation rates are expected to compensate for the shorter growing season, although allocation of photosynthate to cold tolerance or functions other than growth could lead to decoupling of photosynthesis and growth rates (Bazzaz & Grace, 1997; Bosiö et al., 2014; MacArthur, 1984). We thus expect to see a tradeoff between growth and survival in more stressful gardens, as carbon resources from photosynthesis are allocated to tolerating stresses as opposed to growth. Because seedling transplants were grown in natural field conditions that are experiencing ongoing climate change, there may be edaphic differences among sites that can offer more favorable soil resource conditions at some sites relative to others and complicate interpretations of local adaptation in relation to climate.

## Methods

### Reciprocal transplant experimental design

The Adaptation to Climate and Environment (ACE) experiment was designed to test for local adaptation by reciprocally transplanting bur oak seedlings from known families into three garden sites across a latitudinal gradient. The design was based on prior work from Etterson and Shaw (2001), Etterson (2004a, 2004b), Deacon and Cavender-Bares (2015), and Ramirez-Valiente et al. (2018). Acorns from Minnesota, Illinois, and Oklahoma populations were collected in 2018 and 2019 from individual mother trees at least 100 feet apart to minimize the likelihood of shared pollen among trees. Acorns collected from mothers at each site (e.g., park, forest preserve, or natural area) were treated as a single sub-population. Both 2018 and 2019 collections were germinated outdoors in the ground in a common environment in Vallonia, Indiana at the Vallonia State Nursery through the Indiana Department of Natural Resources, grouped by mother tree (see appendix S1: Section S3 supplemental methods for supplemental details on acorn processing). In March 2021, seedlings were lifted from the common nursery, sorted and labelled by mother tree, and sent to be planted in the common gardens in tilled former grasslands in Minnesota (Cedar Creek Ecosystem Science Reserve, East Bethel MN), Illinois (Morton Arboretum, Lisle IL), and Oklahoma (Kessler Atmospheric and Ecological Field Station, Purcell OK). The bare root seedlings were stored in cold storage until the weather was appropriate for planting (Planting times: Oklahoma - March, Illinois - April, Minnesota - May).

In each of the common gardens we planted ten individuals from twenty families from each of the three populations (Minnesota, Illinois, and Oklahoma) 1 m apart in a randomized block design, with at least one individual from each family planted in each block (Fig. 1a). Large herbivores were excluded with fencing around the perimeter of the gardens, and cages were placed around the seedlings to avoid rodent damage. Gardens were weeded or mowed through each growing season and mulched to suppress herbaceous understory growth. The seedlings were irrigated for the first two years after planting to ensure the gardens were established (see Appendix S1: Section S3 supplemental methods for detailed information on garden maintenance).

**Figure 1.**
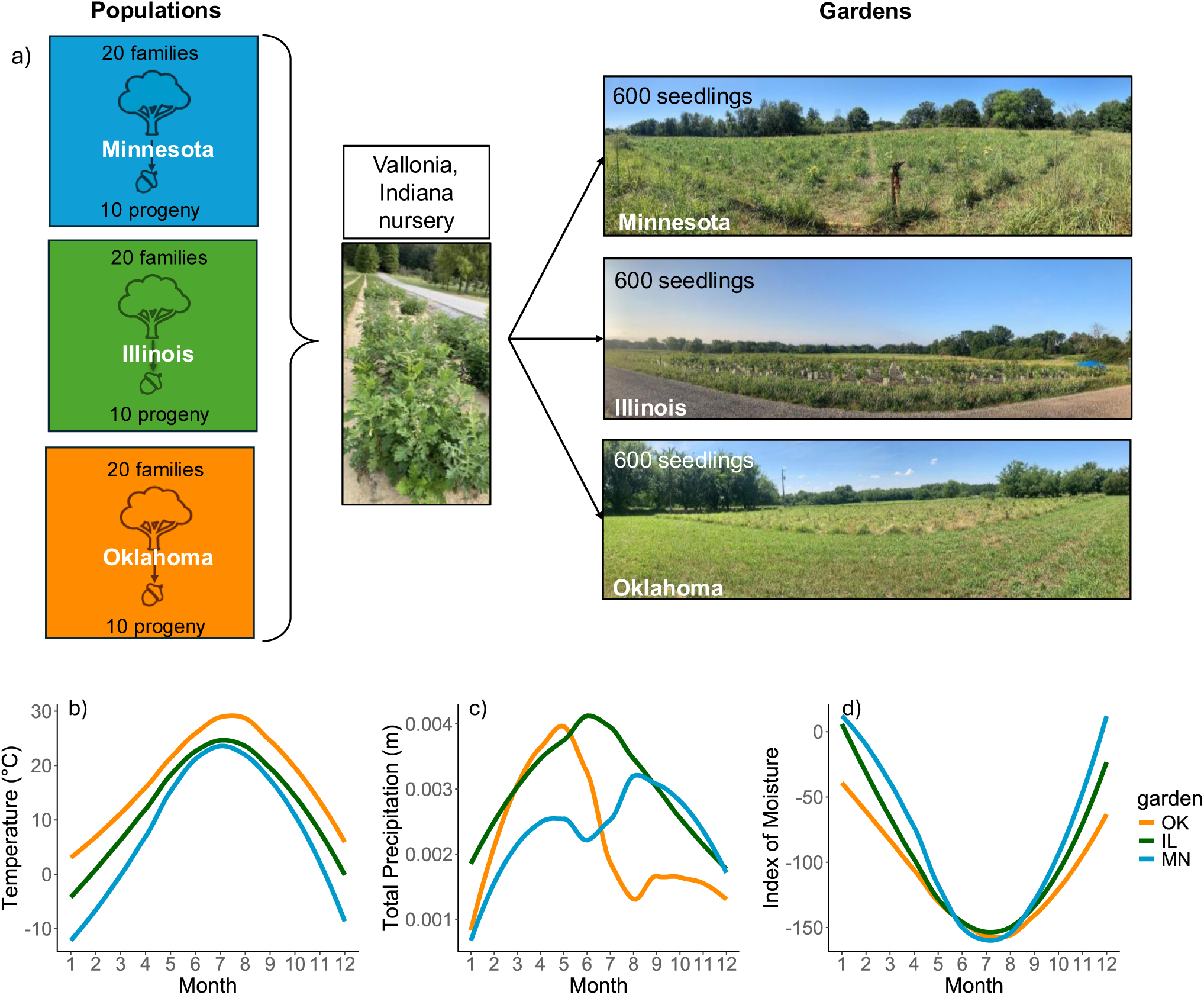
Reciprocal transplant garden design and climate information. In the top panel (a), the experimental design of the Adaptation to Climate and Environment (ACE) experiment is depicted. Three common gardens are located across a latitudinal gradient in Minnesota (Cedar Creek Ecosystem Science Reserve), Illinois (Morton Arboretum), and Oklahoma (Kessler Field Station). In each garden, 600 seedlings were planted representing the three populations from Minnesota (blue), Illinois (green) and Oklahoma (orange). Each population comprised 20 half-sibling maternal families, with half-sibling acorn progeny collected from one known mother tree. Ten progeny from each family were planted in each garden. Photos in each site were taken in summer of 2022 after one year of planting by L.M.S. Rea. In the bottom panel, ERA5 monthly climate data averaged over the years 2021-2023 is shown for temperature (b), total precipitation (c), and the index of moisture (d). Colored lines indicate gardens: Minnesota (blue), Illinois (green) and Oklahoma (orange). Figure created by L.M.S. Rea using icons of “Tree” and “Acorn” from Microsoft Powerpoint.

### Climate data

Monthly temperature data from 1940 to 2023 were acquired from ERA5 data (Hersbach et al., 2023) using garden coordinates and the data extracted from the raster file using QGIS (*QGIS Geographic Information System*, v3.34, 2024). Temperature data for each year were averaged to calculate the mean annual temperature (MAT) and were averaged over the months of May to September to calculate the growing season temperature for each year. To find the change in MAT and growing season temperature since the middle of the 21^st^ century, we calculated the past mean for the years 1940-1960 and the recent mean 2003-2023 then subtracted the past mean from the recent mean for each garden. We used a similar approach to extract precipitation data for the years of 2021-2023, while the garden was planted.

To estimate differences in aridity between the gardens, we calculated an index of moisture (IM) for each of the gardens following Ramírez-Valiente & Cavender-Bares (2017), where the index of moisture is equal to the difference between monthly precipitation and monthly potential evapotranspiration (PET). The IM reflects the monthly water balance for an area and indicates water deficit with negative values and water surplus with positive values. We calculated PET with the Thornthwaite (1948) method, using monthly mean temperature and average day length per month.

### Soil nutrients and texture

To document differences in soil properties among sites, we sampled soil from the center of each garden block to 10 cm depth and conducted tests for soil nutrients (carbon-nitrogen ratios (C:N), total nitrogen content) and soil texture. To measure soil nutrients, we dried the soil and sieved it with a 2 mm mesh sieve to remove large rocks and organic matter. To measure total soil carbon and nitrogen concentration, we ground the soil to a homogenous fine powder and analyzed it with a Costech ECS4010 elemental analyzer (Valencia, California, USA). We approximated the soil texture using the “texture by feel” method (Thien, 1979).

### Survival and growth surveys

To determine seedling performance and fitness, the seedlings were surveyed for survival and size after planting and yearly from 2021 to 2023 at the end of the growing season. The height measured from ground surface to the tip of the highest living bud and basal diameter measured with a caliper 5 cm from the ground were used as metrics of size. Stem volume (V) was estimated using a conoid equation as a proxy for aboveground biomass.

We used conoid volume (V):

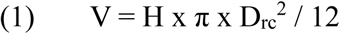

where H is the height of the tree from the root collar to the apical meristem, and D_rc_ is stem diameter at the root collar.

Relative growth rate (RGR) was calculated as

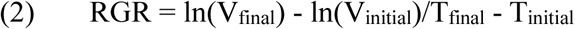

where V is stem volume and T is time in total days. We calculated percent survival of maternal family groups in each garden as the number of trees alive divided by the number of trees originally planted in each family group.

### Phenology

In 2023, to determine growing season length, we tracked spring and fall phenology using protocols established by the National Phenology Network (Denny & Crimmins, 2023). In the spring, approximately weekly, we surveyed the trees in each garden and tracked if the trees were breaking leaf bud. Leaves were considered to be breaking leaf bud if a green leaf tip was visible at the end of the bud. At similar intervals in the fall, we tracked leaves that were browning and/or falling after changing color as senescence. We considered leaves senescing if they had late-season color and were actively falling or had recently fallen from the tree.

We used the budburst and the senescence data to estimate the growing season length of canopy foliage persistence, or how long foliage was present on the trees in a growing season. We calculated this as the difference between the mean initial budburst date and the mean initial senescence date to find the total length of the canopy foliage growing season.

### Gas exchange

To measure photosynthesis rates and stomatal conductance, we conducted measurements of leaf-level gas exchange on every living tree in each of the three gardens using a LI-6800 Portable Photosynthesis System (LI-COR, Lincoln NE, USA) to determine CO_2_ assimilation (A) and intrinsic water use efficiency (iWUE) (see appendix S1: Section S3 supplemental methods for supplemental details on settings and parameters used for measurement). This data was collected in both 2022 (with irrigation) and 2023 (after irrigation ceased) at similar points in the growing season and using the same settings and approach. We calculated iWUE as a ratio of CO_2_ assimilation over stomatal conductance, which is the rate of photosynthetic CO_2_ assimilation against the water loss from open stomata. Because A and iWUE change over the course of the day with stomatal closure, we found the maximum for A, stomatal conductance, and iWUE by excluding measurements after stomatal closure occurred (see Appendix S1: Fig. S3 for cutoff times in each garden for each year).

### Spectral indices

We collected hyperspectral reflectance measurements at the leaf level on every living tree in each garden to add additional, nondestructive metrics of chlorophyll content and nitrogen content. We measured leaf reflectance on the uppermost, fully expanded, undamaged leaf on each tree using a portable field spectrometer (SVC HR-1024i; Spectra Vista) and a leaf clip with an internal light source (LC-RP PRO; Spectra Vista) (see appendix S1: Section S3 supplemental methods for supplemental details on spectral data and processing). Using the spectra, we calculated the chlorophyll normalized difference index (ChlNDI)

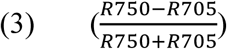

and the normalized difference nitrogen index (NDNI)

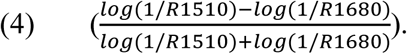

The ChlNDI is positively correlated with chlorophyll a content with minimal influence of chlorophyll fluorescence (Gitelson & Merzlyak, 1994). The NDNI is positively correlated with percent nitrogen of vegetation (Serrano et al., 2002).

### Statistical analysis

#### Aster modeling of fitness

We used aster models to estimate fitness for each population in each garden, using components of fitness expressed early in each trees’ lifecycle, given that the trees were seedlings and not yet reproductively mature at the time of measurement. Aster models estimate fitness integrated over multiple dependent components that contribute to an organism’s fitness. The models avoid issues of nonindependence of normally distributed predictor variables by modeling the dependence of fitness components expressed later in life on earlier fitness components with different statistical distributions (Geyer et al., 2007; Shaw et al., 2008). Aster models thus we can incorporate multiple components of fitness into a unified fitness estimate for each population, rather than having individual predictors of fitness.

We modeled fitness using survival to 2023 and height conditional to survival to the last year of measurements, as our models’ nodes, depicted graphically as:

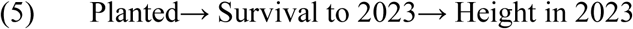

We chose height as the final fitness estimate in our model because it is a useful predictor of eventual fecundity and is dependent on survival (Cornelius, 1994; Gamache & Payette, 2004). Survival to 2023 was modeled with a binomial distribution, and height was modeled with a normal distribution. As predictors in the aster models, we included population (P), garden (G) and the interaction between them, as well as the height at the time of planting (S) as a covariate to account for variation in size that could influence fitness, as:

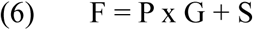

We ran the models using the aster() function in the R package aster (v. 1.3.5. Geyer, 2025). We used the anova() function to compare models with and without the interaction effect to test for significance of the interaction effect. Once we settled on the model (equation 6), we obtained means and standard errors for each population in each garden using the aster predict() function for a dummy dataset with an “average individual” representing each population-garden combination.

#### Statistical testing of performance metrics

We evaluated differences in A_max_, ChlNDI, survival, and growth among and within gardens for all populations using ANOVAs followed by Tukey post-hoc tests with a population by garden interaction effect. The number of individuals in each population in each garden varied due to survival and the number of measurements we were able to obtain (see Appendix S1: Table S1 for number of individuals included in each measure). We also tested relationships between photosynthetic physiology (A_max_ and ChlNDI) with fitness components (survival and growth) using linear models to understand the relationship between physiology and fitness components.

#### Environmental gradients

We conducted principal components analyses (PCA) to understand the environmental gradient captured by the gardens and quantify how the trees’ performance aligns with the environmental gradient. As climate variables, we calculated minimum and maximum monthly average temperatures, the average growing season precipitation (as monthly averages of May-September precipitation), and minimum index of moisture (IM) (as the minimum IM in May-September) for each garden. For soil variables, we included the percent nitrogen, percent carbon, and percent C:N ratio for each block in each garden. We centered and scaled the variables and used the prcomp() function to conduct the PCA. We extracted the PCA scores for PC1 and PC2 and evaluated their associations with performance metrics including A_max_, ChlNDI, NDNI, survival, growth, and fitness using linear models. We used generalized linear models for analysis of PCA scores with survival using the function glm() and the “logit” link function, given that the survival data is binomial.

## Results

### Site climate differences

The ERA5 dataset showed that the average monthly temperature decreased latitudinally, with the Minnesota garden having the lowest average temperatures each month and Oklahoma having the highest (Fig. 1b). Notably, the Oklahoma garden never had a monthly average below freezing (0 °C), whereas the Minnesota and Illinois gardens did in the winter months. We found that since 1940, temperatures have increased at each of the garden sites. Mean annual temperature (MAT) has increased by over 1°C in each garden from averaged 20-year periods of 1940-1960 to 2003-2023, and the amount of temperature change increased with latitude, with the greatest change in MAT occurring in the northernmost garden, Minnesota, and the lowest change in temperature in the southernmost garden, Oklahoma (Table S2). Over the same period, the average growing season temperature has also increased. Like MAT, the greatest change in growing season temperature was in Minnesota. However, unlike MAT, the smallest bchange in growing season temperature was in Illinois, the middle garden.

The gardens also differed in precipitation and aridity over the year. The Oklahoma garden peaked in precipitation in the springtime around April and May (Fig. 1c). The Minnesota garden also had a small peak in precipitation at that time, however the highest monthly precipitation occurred in late summer for the Minnesota garden, around August and September. Precipitation in the Illinois garden peaked mid-summer, in June and July, and was the highest precipitation of the year among the three gardens. The Illinois garden was also the least arid of the three gardens during the growing season, though the index of moisture was negative for all the gardens, indicating a water deficit during the growing season (Fig. 1d). Notably, the index of moisture was negative year-round for the Oklahoma garden, suggesting it was in an atmospheric water deficit for the entirety of the year while the gardens were planted, though some of the water stress was presumably alleviated by irrigation in the first two years of growth.

### Fitness components

We assessed two main components of fitness: survival and plant size. Survival rates of family groups were most disparate in the Oklahoma garden, where survival of the Minnesota population (MN) was significantly lower than the Illinois (IL) and Oklahoma (OK) populations (*F*(2, 181) = 8, p = 0.0004). The higher survival of OK compared to MN is in line with local vs. foreign local adaptation (Fig 2a). Garden had a strong effect on survival percent *F*(2, 181) = 34, p = 2.88e-13). Survival for MN and IL was highest overall in the Illinois garden. Survival was not significantly different across populations in the Illinois and Minnesota gardens. There was a significant population × garden interaction for survival, as survival differed across gardens, particularly for the IL and MN populations (*F*(4, 181) = 5.5, p = 0.0003).

**Figure 2.**
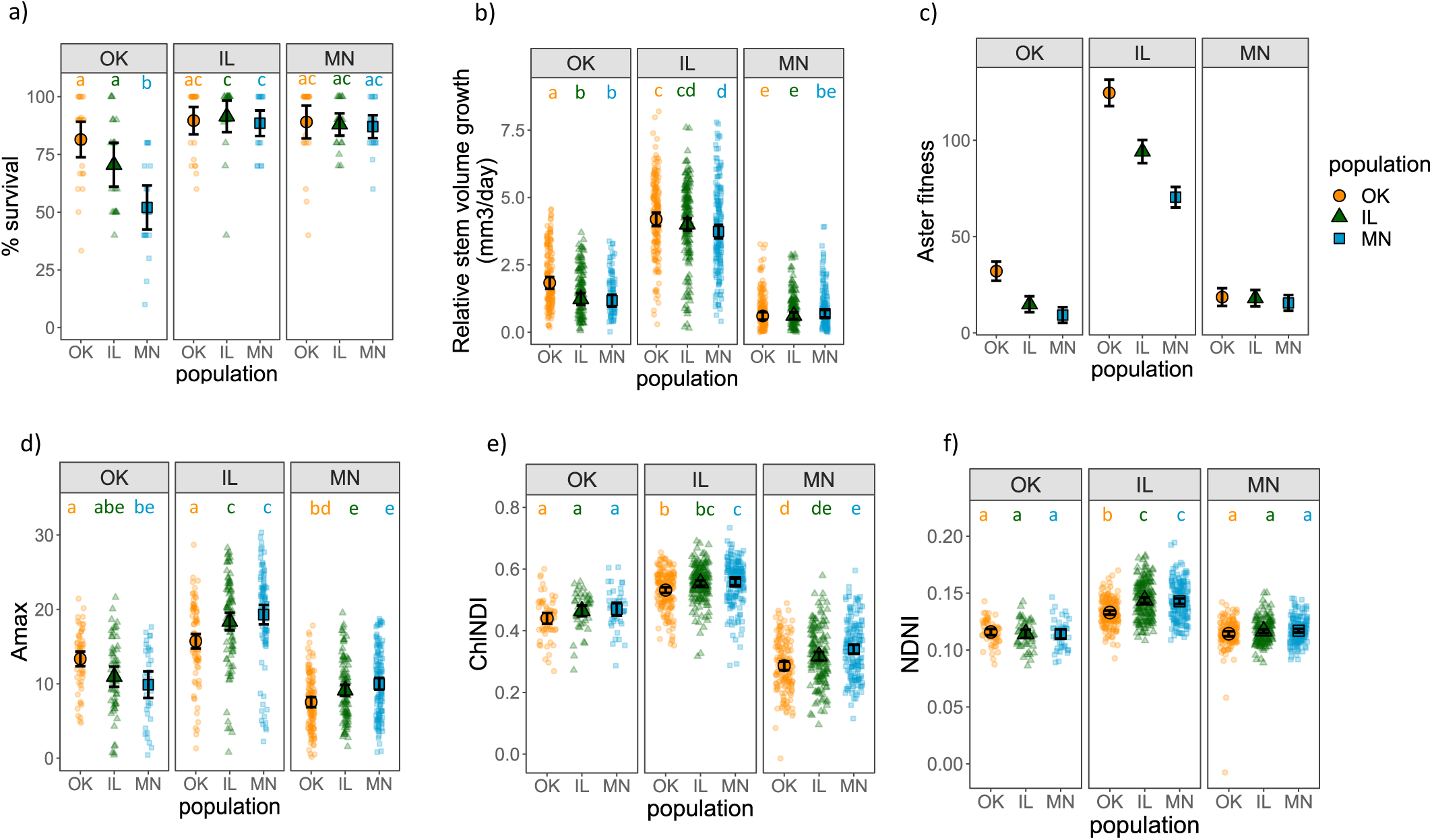
Fitness, fitness components, and physiology in each garden. Each panel represents the common garden located in Oklahoma (OK), Illinois (IL), or Minnesota (MN). Progeny from each source population are represented by orange circles for OK, green triangles for IL, and blue squares for MN, with larger points indicating population means. Distinct letters denote statistical significance (p<0.05) for each population group. Percent survival is represented by a point for each maternal family (a), and was highest for the local OK population in OK, but was similar among all populations in the northern gardens. The relative growth rate of stem volume for individuals (b) was also highest for the local population in OK, with the lowest growth rates occurring in the MN population. Aster expected fitness (c) is represented by a point for each population, and was highest in the Illinois garden. Aster fitness significance was determined by omnibus tests of population*garden interactions. Maximum CO_2_ assimilation (A_max_) (d) of the local populations was highest for the OK and MN garden, but overall the IL garden had the highest A_max_. Spectral chlorophyll normalized difference index (ChlNDI) (e) was also higher overall in the IL garden for all populations, though the local MN population was highest in the MN garden. Normalized difference of the nitrogen index (NDNI) (f) indicates that leaf nitrogen was highest in the Illinois garden.

Stem volume combines the effects of diameter and height growth. There was a significant effect of population on relative growth rate of stem volume (RGR) *F*(2, 1523) = 5.37, *p* = 0.0047. The OK population had the highest stem volume RGR in the Oklahoma garden but was similar to the IL population in the Illinois garden (Fig. 2b). Garden also had a strong effect on stem volume RGR, (*F*(2, 1523) = 704.46, *p* < 0.0001). In the Minnesota garden, the stem RGR was similar among all populations. Stem volume RGR was highest in the Illinois garden for all populations. In addition, the population × garden interaction was significant for RGR, (*F*(4, 1523) = 4.34, *p* = 0.0017).

### Fitness

We found significant (χ²(65) = 125.08, p = 0.00001) interaction effects of population × garden on fitness using omnibus tests. Block was not a significant predictor of fitness. Our model showed that aster fitness was highest in the Illinois garden for all populations, with the OK population having the highest fitness, in line with our hypothesis of climate change having shifted ranges northward. (Fig. 2c). In the Oklahoma garden, the OK population had higher fitness than the northern populations, as expected with local vs. foreign local adaptation. Fitness in the Minnesota garden was about the same for all populations.

### Physiology

We evaluated photosynthetic responses in each garden. There was a significant population effect for the maximum CO_2_ assimilation (A_max_), (*F*(2, 804) = 8.8, p = 0.0002). A_max_ was highest for the MN and IL populations in the Illinois garden (Fig. 2d). Garden had a significant strong effect on A_max_, (*F*(2, 804) = 274, p< 2e-16). The MN and IL populations had similar A_max_ rates to each other in all gardens and were lower in the Oklahoma garden than in the northern gardens, as we hypothesized given the aridity stress in the Oklahoma garden. There was a significant population × garden effect, (*F*(4, 804) = 9.7, p = 1.2e-7). The OK population had lower A_max_ rates in both the Minnesota and Illinois gardens compared to the northern populations but had higher A_max_ rates in the Oklahoma garden compared to the MN population.

We found significant effects of ChlNDI with population (*F*(2, 1168) = 22.2, p= 3.3e-10) and a strong effect with garden (*F*(2, 1168) = 1209, p< 2e-16), but no significant interaction effect. The OK population also had lower chlorophyll content (as detected by the chlorophyll normalized difference spectral index (ChlNDI)) in the Illinois and Minnesota gardens compared to the MN population (Fig. 2e). The IL population ChlNDI values were generally in between the OK and MN populations. There was a significant effect of spectral normalized difference nitrogen index (NDNI) with population, (*F*(2, 1168) = 25, p= 2.3e-11) and a strong effect with garden (*F*(2, 1168) = 470, p< 2e-16), as well as a population × garden effect (*F*(4, 1168) = 8.1, p< 2e-6). We found that NDNI was highest for all populations in the Illinois garden, indicating higher leaf nitrogen content (Fig. 2f). The OK population had lower leaf nitrogen content than the northern populations in the Illinois garden. The NDNI was similar for all populations in the Oklahoma and Minnesota gardens. Data from 2022 was significantly different from 2023 data but indicated similar though weaker trends, which may be an effect of irrigation or ontogeny. We focus on 2023 data here as the effects are stronger and indicate responses in natural conditions without irrigation (see Appendix S1: Fig. S4 for 2022 results).

### Correlations of survival, growth, and physiology

Stem volume growth was significantly (p<0.05) positively correlated with survival for all populations in the Illinois garden (OK t=2.4, df=22, p=0.02; IL t=4.3, df=17, p=0.0004; MN t=2.1, df=18, p=0.47), but the relationship was not significant in the Minnesota or Oklahoma gardens (Fig. 3a). We found significant (p<0.05) positive relationships of A_max_ with stem volume growth for the OK and IL populations in the Oklahoma (OK t=4.1, df=60, p=0.0001; IL t=3.7, df=50, p=0.0005), Illinois (OK t=6.7, df=98, p=1.2e-9; IL t=6.8, df=94, p=7.9e-10), and Minnesota (OK t=2.5, df=120, p=0.012; IL t=3.7, df=109, p=0.0003) gardens, and for the MN population in the Illinois garden (t=6.2, df=94, p=1.5e-8) (Fig. 3b). We also found significant (p<0.05) positive correlations of ChlNDI with stem volume growth in the Illinois garden for all populations (OK t=6.9, df=176, p=8.8e-11; IL t=5.1, df=180, p=7.4e-7; MN t=6.4, df=178, p=1.6e-9), and for the IL and MN populations in the Minnesota garden (IL t=4.4, df=159, p=1.6e-5; MN t=3.4, df=162, p=0.00095) (Fig. 3c). We did not find significant relationships with survival for A_max_ or ChlNDI (not shown).

**Figure 3.**
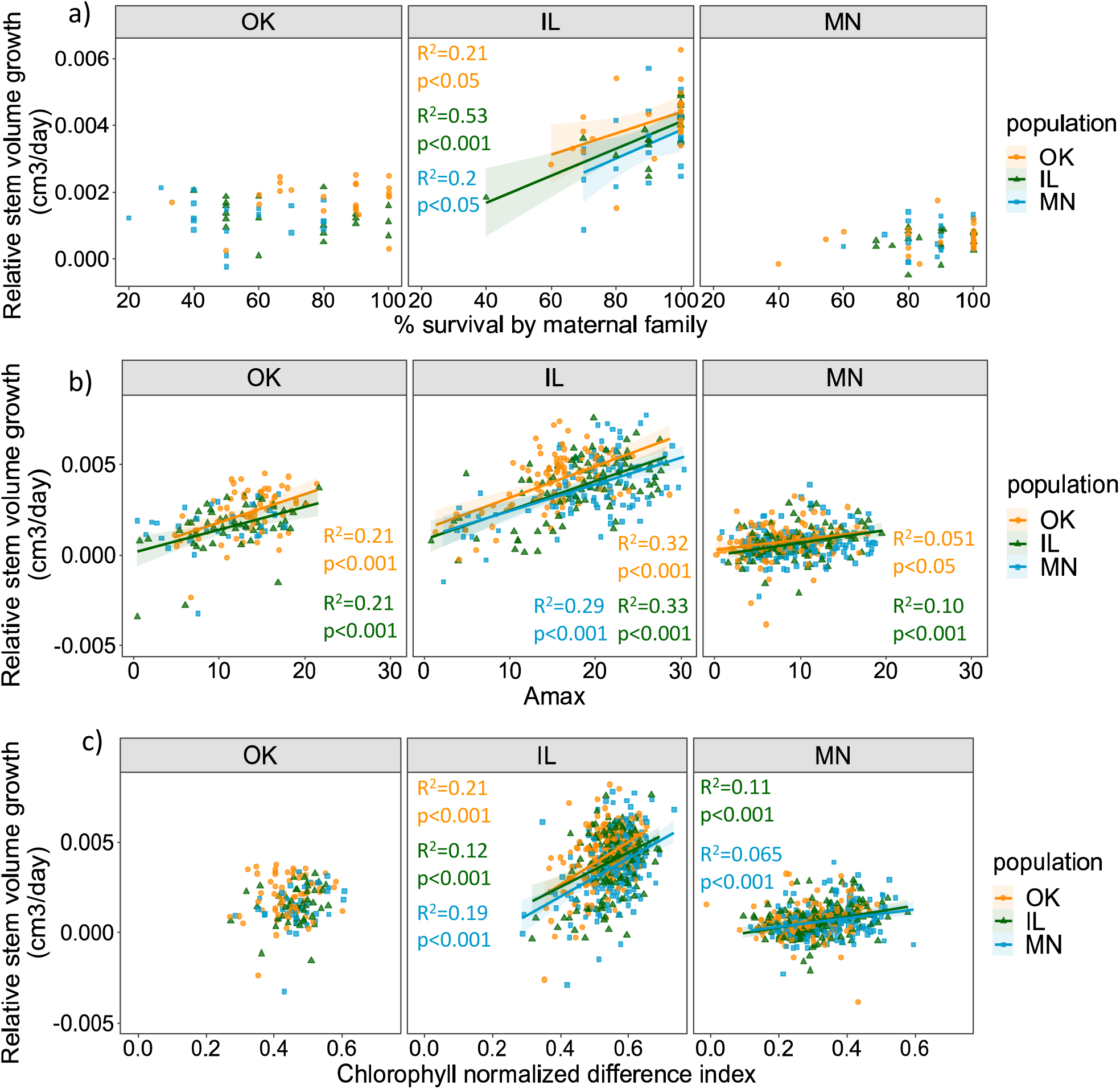
Comparisons of growth, survival, and physiology from the 2023 data collection period to understand shifts in carbon allocation. Each panel represents a common garden located in Oklahoma (OK), Illinois (IL), or Minnesota (MN). The populations are represented by an orange circle for OK, a green triangle for IL, and a blue square for MN, with significant trendlines (with standard errors) in matching colors. a) Percent survival by maternal family is significantly positively correlated with relative stem volume growth for all populations in the Illinois garden, but not in the OK or MN gardens. Maximum photosynthetic rate (A_max_) has significant relationships with growth (b) for OK and IL populations in all gardens, and for the MN population in the IL garden. Chlorophyll normalized difference index (ChlNDI) was significantly positively correlated with relative growth rate (c) in IL for all populations, and was significantly correlated with growth in the MN garden for the IL and MN populations.

### Soil quality

Soils in the Oklahoma and Illinois gardens were very similar in carbon (C) and nitrogen (N) concentrations, though concentrations in Oklahoma were a bit higher on average (Table 1). Soils in the Minnesota garden had low percent nitrogen and percent carbon. Minnesota had the highest soil C:N, followed by the Illinois and then the Oklahoma gardens. The soils also differed texturally. Oklahoma garden soils were classified as silty clay loam, Illinois as sandy clay loam, and Minnesota as sandy loam.

**Table 1.** Soil properties in each of the three gardens.

|  | % | % |  |  |
| --- | --- | --- | --- | --- |
| Garden | Nitrogen | Carbon | C:N | Soil texture |
| MN | 0.06 | 1.04 | 21.32 | Sandy loam |
| IL | 0.25 | 3.54 | 14.12 | Sandy clay loam |
| OK | 0.30 | 3.33 | 11.21 | Silty clay loam |

### Environmental gradients across gardens

The gardens fall across two main PCA axes that together account for 97.9% of the total environmental variance (Fig. 4). PC1 accounts for 57.5% of the variance and represents the gradient of temperatures and soil nutrients, with higher values corresponding to higher temperatures and nutrients. PC2 captures 40.4% of the variance and has higher values corresponding to higher precipitation and higher IM (meaning less aridity). The Oklahoma garden falls on the right side of PC1 and the bottom of PC2, indicating that it has high temperatures and nutrients, but low precipitation and high aridity. The Minnesota garden similarly falls on the bottom of the PC2 axis, but is to the left of the PC1 axis, indicating it is a dry environment, with lower temperatures and lower nutrients. The Illinois garden lies on the top end of the PC2 axis, indicating high precipitation and low aridity, but falls in the middle of PC1 putting it at intermediate soil nutrients and temperatures.

**Figure 4.**
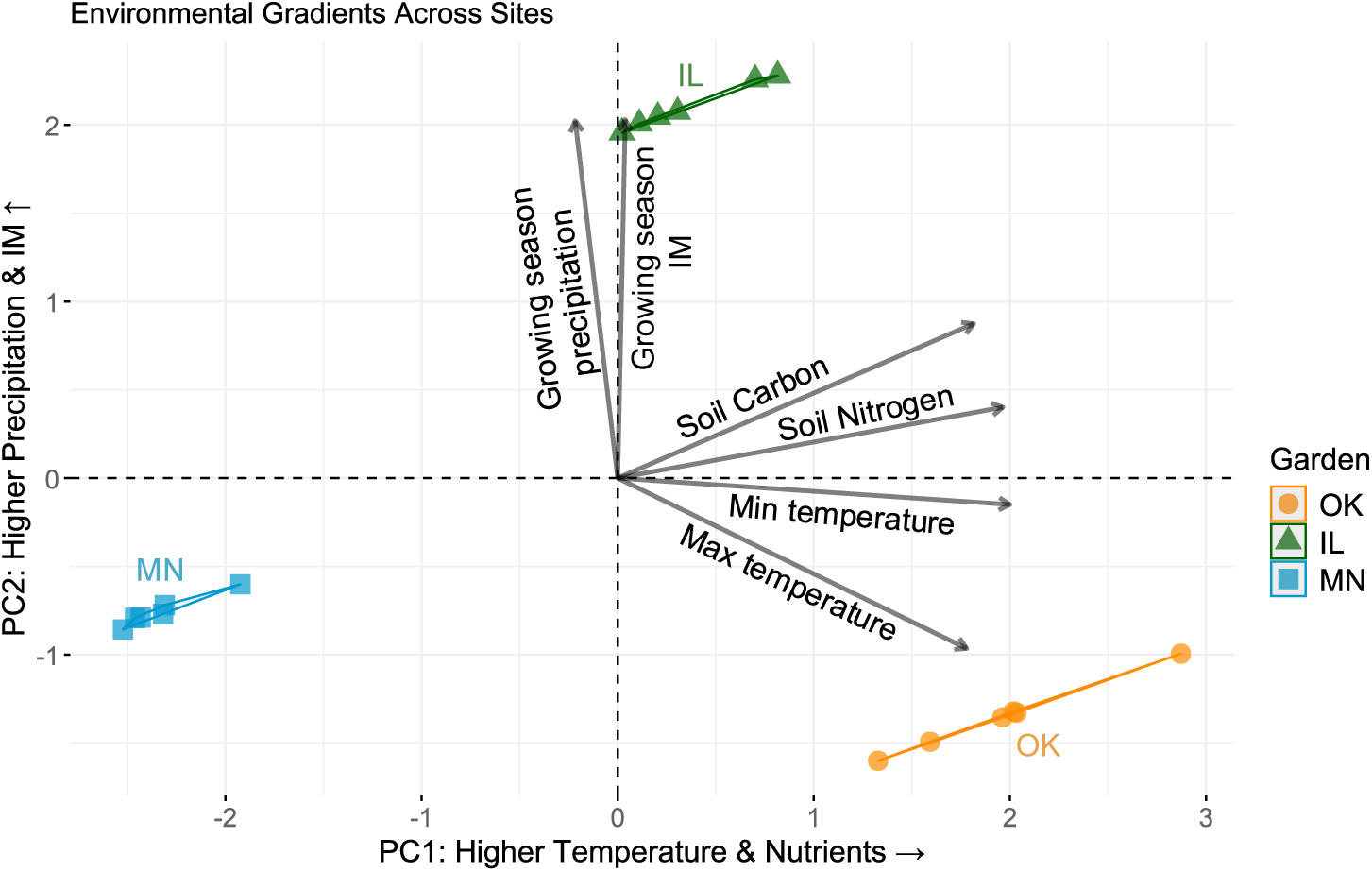
PCA biplot depicting environmental gradients across gardens, with PC1 plotted as the x axis and PC2 plotted on the y axis. Climate and soil conditions including growing season precipitation, growing season index of moisture (IM), soil carbon, soil nitrogen, minimum (min) temperature, and maximum (max) temperature are indicated by labeled arrows, with the direction indicating where they fall along the PCA axes. The estimates for sampled garden blocks in Oklahoma (OK), Illinois (IL), and Minnesota (MN) gardens are plotted as points based on where their environmental conditions fall along the gradient between PC1 and PC2. The points are connected with a convex hull polygon that bounds the observed values for each garden. This PCA indicates that temperature and soil nutrients increase with PC1 and precipitation and moisture increase with PC2.

Regressions of performance metrics on PCA axes indicate that PC2 is a consistently strong predictor of performance, indicating that less arid climates lead to better performance and survival (Table 2). PC2 was a particularly strong predictor for A_max_, fitness, and survival. PC1 was positively associated with all traits except survival, indicating that environments with hotter temperatures and higher nutrients lead to lower survival.

**Table 2.** Results of regressions of PCA axes with performance metrics of photosynthetic rate (A_max_), fitness, chlorophyll normalized difference spectral index (ChlNDI), leaf nitrogen normalized difference index (NDNI), survival, and relative growth rate. R^2^ values are reported for all but survival, where the McFadden pseudo - R^2^ is reported to compare the log likelihood of the full GLM model to the log likelihood of the null model.

| Response Variable | Model Type | PC1 Effect | PC2 Effect | Degrees of freedom | $R^2$ / Pseudo-$R^2$ |
| --- | --- | --- | --- | --- | --- |
| $A_{max}$ | Linear | +0.72<br>($p=4.5e-13$,<br>$t=7.4$) | +2.23<br>($p<2e-16$,<br>$t=17.7$) | 810 | $R^2=0.376$ |
| **Fitness** | Linear | **+4.20**<br>(p< 2e-16,<br>t=12.4) | **+24.37**<br>(p< 2e-16,<br>t=60.2) | 1788 | R<sup>2</sup>=0.679 |
| **ChlNDI** | Linear | **+0.0333**<br>(p< 2e-16,<br>t=22) | **+0.0473**<br>(p< 2e-16,<br>t=27.8) | 1174 | R<sup>2</sup>=0.644 |
| **NDNI** | Linear | **+0.00052**<br>(p=0.049,<br>t=2) | **+0.00779**<br>(p< 2e-16,<br>t=26.4) | 1174 | R<sup>2</sup>=0.433 |
| **Survival** | Logistic | **-0.213**<br>(p=6.5e-10,<br>z=-6.12) | **+0.310**<br>(p=3.9e-11,<br>z=6.6) | 1798 | McFadden=0.06 |
| **Relative**<br>**Growth**<br>**Rate** | Linear | **+0.00023**<br>(p< 2e-16,<br>t=13.9) | **+0.00081**<br>(p< 2e-16,<br>t=32.4) | 1529 | R<sup>2</sup>=0.468 |

### Phenology and growing season

Budburst was recorded earliest in the Oklahoma garden and latest in the Minnesota garden, on average. We found significant effects of budburst with population *F*(2, 984) = 78.6, p<2.2e-16, garden *F*(2, 984) = 508.8, p<2.2e-16, and a significant interaction of population × garden *F*(4, 984) = 16.3, p=6.2e-13. Of the three populations, the OK population tended to break bud earliest on average, except in the Minnesota garden where all populations broke bud at similar times (Fig. 5a). We found significant effects of fall senescence with population *F*(2, 984) = 33, p<2.2e-16, garden *F*(2, 984) = 218.5, p<2.2e-16, and a small but significant interaction of population × garden *F*(4, 984) = 2.9, p=0.02. The Minnesota garden had the earliest date of senescence, whereas the Oklahoma garden was the latest (Fig. 5b). The OK population also tended to senesce later than the northern populations, particularly in the Illinois garden. The Oklahoma garden had the longest growing season length for all three populations, whereas Minnesota was the shortest (Fig. 5c).

**Figure 5.**
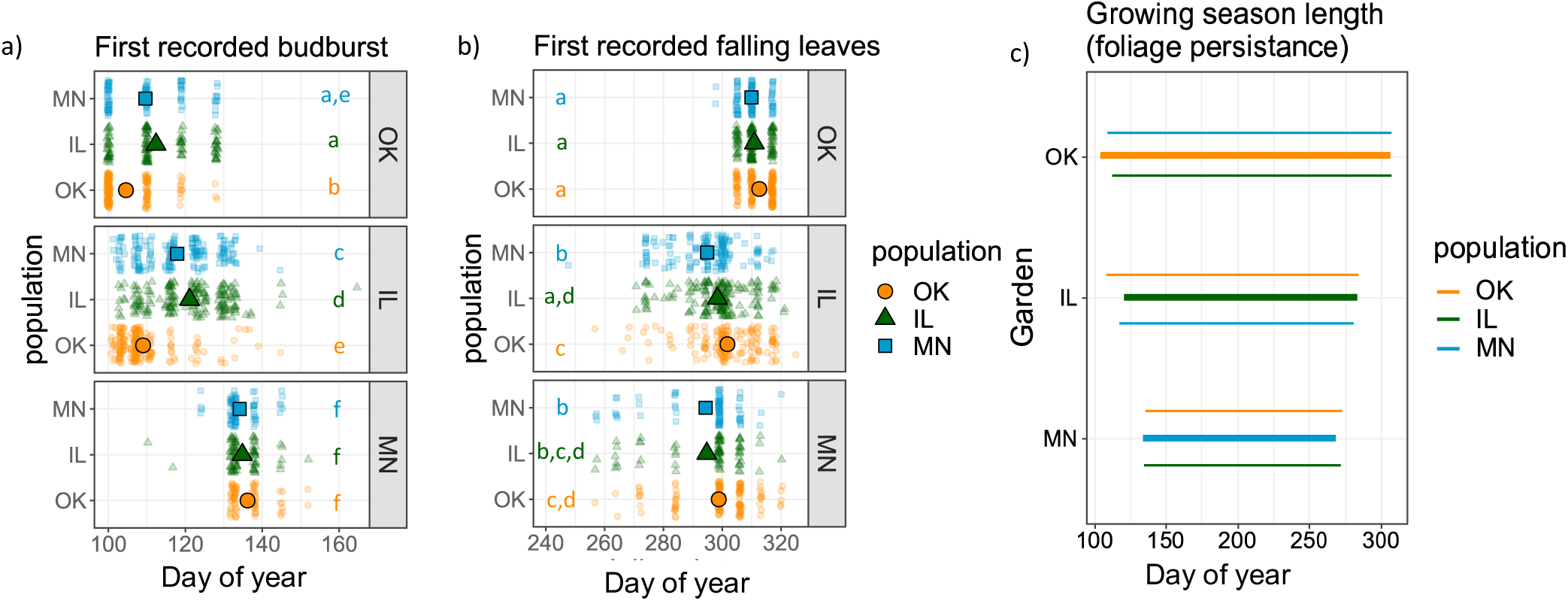
Phenology and growing season length in each of the gardens with data collected in 2023. Each panel represents a common garden located in Oklahoma (OK), Illinois (IL), or Minnesota (MN). The populations are represented by an orange circle for OK, a green triangle for IL, and a blue square for MN. Budburst (a) occurs earlier in the OK and IL garden for all populations than in the MN garden, and senescence (b) occurs latest in the OK garden. The growing season length (c) is shortest in the MN garden and longest in the OK garden.

## Discussion

### Evidence for local adaptation

We established the ACE (Adaptation to Climate and Environment) experiment, a reciprocal transplant experiment along a latitudinal gradient, to test for local adaptation and adaptive differentiation among populations of *Quercus macrocarpa* from contrasting climates and environments. For all populations, we expected that local adaptation would be demonstrated by higher fitness of the local population relative to other populations (local vs foreign) and higher fitness of each population in its home location relative to its fitness in other locations (home vs. away). We found evidence supporting local adaptation based on the local vs. foreign comparison in the Oklahoma garden with the hottest climate. The OK source population had higher survival, growth, and fitness in the Oklahoma garden, compared to the other populations, which supports local vs. foreign adaptation. Under the 2 °C warming expected (IPCC, 2023) with climate change, the OK population may be better suited for the emerging warmer climates with potential to expand northward. These results can inform ongoing efforts to manage forests and identify seed collection zones for transferring seeds among regions by demonstrating that the southernmost population is adapted to warmer climates. Populations adapted to warmer climates may be suitable for reforestation efforts aimed at improving forest resilience to climate change or transitioning ecosystems to novel climates (Palik et al., 2022).

Despite evidence of local adaptation in the Oklahoma garden, our results show that all populations expressed highest fitness in the Illinois garden. Consequently, we did not find evidence for local adaptation based on home vs. away comparisons. This effect may be due in part to differences in management—the gardens were watered on differing schedules (Appendix S1: Section S3 supplemental methods), and the Illinois garden was also weeded more intensively than the other two gardens—but the effect not entirely or, we believe, even primarily: the differences between the three gardens in precipitation, extremes of temperature and precipitation, and soils are profound (Figures 1b–d, 4). We also did not find clear statistical evidence for local vs. foreign local adaptation in the two northern populations in their respective gardens after three years of growth. However, these patterns of local adaptation could still emerge through time or could be obscured by maternal effects. The OK population had higher absolute growth rate, which could be due in part to its larger acorn size (Appendix S1: Table S3), with larger seed size supporting growth and survival in its early years (Bischoff et al., 2006; Deacon & Cavender-Bares, 2015). We were not able to measure all critical ontogenetic stages and lifetime components of fitness, including germination and fecundity. Reproductively mature life stages could still reveal evidence of local adaptation not evident at the seedling life stage (Wadgymar et al., 2022). Nevertheless, the patterns of maladaptation to warmer climates we observe in northern populations could also increase in later life stages (Goetz et al., 2024, 2025). It is critical to study seedling survival and function to understand the future of tree populations given that seedlings often have a narrower climatic niche and are more vulnerable compared to adult trees, but their establishment is critical to the regeneration and persistence of tree populations (Davis et al., 1998; Dobrowski et al., 2015; N. Williams, 2021).

Our alternative hypothesis for the observation of high performance outside of populations’ local gardens was that ongoing climate change has shifted suitable ranges of all populations northward, leading southern populations to be better adapted to northern gardens. However, while we saw that the OK population did perform better in the Illinois garden, which could be a consequence of both edaphic and climatic factors, we did not see evidence that the IL population performed better in the Minnesota garden compared to in its home garden, nor did it outperform the local MN population in Minnesota. This result indicates that there may be additional conditions or stressors in the Minnesota garden, such as freezing, nutrient deficiencies, and/or insect herbivory, that influence performance of non-local populations.

### Variation in season length helps explain carbon assimilation and growth patterns

Higher survival, growth rates, and fitness in the Illinois garden for all populations were likely a consequence of productive soil conditions and mild climate relative to more stressful environments (the cold of Minnesota, the aridity of Oklahoma) found closer to the margins of *Q. macrocarpa’s* range. Aridity stress in the Oklahoma garden led to lower photosynthetic rates in northern populations, as evidenced by the association between moisture levels and A_max_ rates (Table 2). The decline in assimilation rates in seedlings from the IL and MN populations in the hotter and drier Oklahoma climate suggests that these populations may experience physiological stress leading to future declines in survival and fitness in their current habitats. In addition, the OK population had the highest maximum carbon assimilation rate (A_max_) of the three populations in the Oklahoma garden. The lower A_max_ rate of the northern populations in the Oklahoma garden indicates they were stressed in the hotter climate. Drought is known to reduce stomatal conductance, which limits carbon assimilation (Boyer, 1976), and the Oklahoma garden was also the most arid. Previous findings from oaks have shown that A_max_ is reduced when plants are exposed to chronic drought (Gimeno et al., 2008).

Differences in growing season lengths along the latitudinal gradient also played a role in the populations’ photosynthetic and growth responses. Frost damage may be causing dieback of stems (negative growth) of the OK population in the Minnesota garden, likely due to later senescence, consistent with previous work in another North American temperate oak species (*Q. rubra*) that implicates autumn frost damage at northern range limits growth of trees in southern seed sources (Lindback et al., 2023). The OK population broke bud earlier and senesced later in the Oklahoma and Illinois gardens relative to other populations, consistent with findings in other oak common gardens (Papper & Ackerly, 2021; Wright et al., 2021) and supporting our expectations of populations from lower latitudes being adapted to longer growing seasons. The Oklahoma population tended to maintain a relatively low rate of carbon assimilation across gardens; compared to the other populations in the northern gardens, it had the lowest carbon assimilation rate. This is in accordance with expectations for slow life history strategies expectations, where leaves with longer life spans are theoretically expected and empirically demonstrated to have slower rates of photosynthesis (Chabot & Hicks, 1982; Wright et al., 2004). On the other hand, the northern populations that are adapted to shorter growing seasons tended to have higher A_max_ rates than the Oklahoma population in the northern gardens. This trend of higher photosynthetic rates is consistent with findings in a *Betula pendula* providence trial in which northern populations had greater net photosynthesis (Tenkanen et al., 2021), and has implications for the growth and survival of northern populations as climate change extends growing season length. These stressors of heat (Oklahoma) and frost (Minnesota) in the gardens at the ends of our latitudinal gradient likely explain the high performance of seedlings in the milder Illinois garden.

Stressful conditions can also shift carbon allocation away from growth toward drought stress responses and defense (Wang et al., 2024). We found correlations between growth and survival in the Illinois garden, in which all populations expressed highest fitness, likely due to its relatively mild climate and loamy soils. Previous work in the European pedunculate oak, *Quercus robur,* indicates that differences in soil conditions and water availability across sites with similar climate regimes lead to variation in productivity (Buras et al., 2020). Growth and survival were decoupled in both the Oklahoma and Minnesota gardens, which may be a consequence of resource allocation trade-offs under increased stress (heat in Oklahoma, cold and low nutrient availability in Minnesota, and aridity in both) that improves survival at the expense of growth rates (Bazzaz et al., 1987; Willi & Van Buskirk, 2022). In addition, despite having higher soil nitrogen levels, the fine textured silty-clay soils in Oklahoma may be less conducive to nutrient access.

Carbon assimilation and light energy acquisition through photosynthesis were important for growth, as evidenced by the positive correlations between growth and A_max_ and leaf chlorophyll content, particularly in Illinois. In fertile sites, relative growth rates and photosynthetic rates tend to be higher, as was the case in the Illinois garden, where conditions were favorable (Lambers & Poorter, 1992). In the Illinois garden, the seedlings were able to allocate most of their carbon toward aboveground growth, which in turn improved survival. On the other hand, in the less favorable, more arid, conditions of Minnesota and Oklahoma, survival likely depended on allocating carbon elsewhere, such as drought adaptation, root growth, or defense against herbivory (Coley et al., 1985; Poorter et al., 2012). Selecting populations that are able to shift allocation of carbon to improve survival will likely be important given the expected increased intensity of drought and pests with climate change.

### Multiple environmental factors vary across gardens

An outdoor common garden experiment such as ours, with unmanipulated soils, cannot isolate climate as the sole factor that varies among sites; and it may well turn out that the seedlings in the Illinois garden perform best due to non-climatic factors. Edaphic factors, including unmeasured factors such as the microbial environment, vary substantially among experimental replicates (in three states). We found that the environmental conditions in gardens fell along a gradient of soil nutrients and temperature and a gradient of aridity and precipitation. The Illinois garden had high soil nitrogen and organic matter content. Leaves from plants across populations in the Illinois garden had higher spectrally detected nitrogen content (NDNI).

Nitrogen is a major component of photosynthetic chemistry, including RuBisCo and chlorophyll, and is consequently correlated with higher photosynthetic rates (Ellis, 1979; Evans, 1989; Luo et al., 2021). Total soil nitrogen concentration was similar between the Illinois and Oklahoma gardens, but the less arid and slightly cooler climate in the Illinois garden combined with the high soil nitrogen and soil organic matter of the loamy soil created an environment that is conducive to bur oak carbon assimilation and fitness. We also note that of the three gardens, Illinois had the least change in growing season temperature since 1940 (Appendix S1 Table S2). The fertile soil and relatively stable, mild climate of the region near the Illinois garden may provide important habitat for bur oak populations to grow with ongoing climate change.

Controlled environment experiments that isolate climate as the driving variable for performance changes in bur oaks are an important next step in predicting response of bur oak populations to climate change, following experiments such as (Browne et al., 2019b; Sáenz-Romero et al., 2017), which have tested climate response in other oak species in mediterranean climates, which have found evidence of adaptational lag and negative responses of trees to warmer climates.

## Conclusion

Findings from the ACE reciprocal transplant experiment provide context for seed transfer aimed at improving resilience of northern tree populations in the face of climate change. Our results indicate that northern bur oak populations may suffer increasing mortality in the face of climate change. We found evidence that the northern population was poorly adapted to hotter conditions in the south, and populations originating from warmer southern climates perform well even when they are planted northward into Illinois. These findings indicate that southern locations may provide valuable seed sources to augment existing genetic variation in northern regions in the face of ongoing climatic warming. Consideration of current climate and environmental conditions alongside future climate conditions are important in decisions to move locally adapted populations. It may be beneficial to ensure seeds are sourced from locations with varied climates to accommodate fluctuations in climate. It is also worth noting that the expected change in climate may be within the tolerance limits of existing populations in the north. Mild climate warming akin to that experienced by the MN populations in the Illinois garden may even improve performance and fitness in the case of *Q. macrocarpa.* However, the poor performance of the MN population in the warmest garden may need to be considered in long-term forest planning, given the rapid rates of warming in northern latitudes, and may require seed transfer from southern seed sources to supplement and maintain northern bur oak populations adapted to warmer climates. We did not test performance under competition with other species nor explicitly vary edaphic environments across climates, however, the performance of oaks across contrasting edaphic environments in competition with other tree species is an important area of investigation for future studies.

## Acknowledgements

We would like to thank those without whom the work presented here could not have been done. We thank Kris Bachtell, Cathy Bechtoldt, Brett Fredericksen, Marlene Hahn, Troy Mielke, Maria Park, Senna Robeson, Allison Scott, Nick Stoynoff, the Cedar Creek Ecosystem Science Reserve, Morton Arboretum Facilities and Natural Resources staff, and The Kessler Atmospheric and Ecological Field Station staff for help establishing the common garden. We thank the Indiana Department of Natural Resources Vallonia Nursery for the use of their facilities and help with harvesting seedlings. Field assistants including Iris Cessna, Isaiah Clark, Allison Collins, Nick Duncan, Rachel Gowett, Jennifer Hamann, Leah Hill, Keion Mackey, and Morton Arboretum volunteers assisted with data collection. Ruth Shaw provided instruction on aster modelling and provided edits on an earlier version of the manuscript. We acknowledge funding from ASCEND Biological Integration Institute (NSF DBI 2021898), the National Science Foundation Dimensions of Biodiversity grant (NSF DEB 2129312 (UMN), 2129281 (MOR), and 2129236 (UO)), the Morton Arboretum Center for Tree Science, and Cedar Creek Ecosystem Science Reserve Long Term Ecological Research Program through NSF DEB 1831944.

## Author Contribution Statement

All authors contributed intellectually to the study. JCB designed the ACE experiment and managed its implementation. JCB, ALH, HRM managed the collaboration and secured funding for the study. LMSR led data collection, analysis and interpretation, and writing of the manuscript. JCB led nursery planting. CL, LW, and MG led planting efforts in the gardens. LO, RM, MG, LW, and CL collected data. JCB, ALH and HRM edited early versions of the manuscript. All authors provided critical revision of the article and provided final approval of the version to be published.

