## Appendix S1 for "Genetically based variation in fitness and carbon assimilation among bur oak populations"

**Journal name:** Ecology

**Manuscript type:** Article

**Manuscript title:** Genetically based variation in fitness and carbon assimilation among bur oak populations

**Author names:** Lucy M.S. Rea^1^, Laura Ostrowsky^1,2^, Rebekah Mohn^3^, Mira Garner^3,4^, Lindsey Worcester^3^, Cathleen Lapadat^1^, Heather R. McCarthy^5^, Andrew L. Hipp^3^, Jeannine Cavender Bares^1,2*^

**Affiliations:**

^1^Department of Ecology, Evolution, and Behavior, College of Biological Sciences, University of Minnesota, 140 Gortner Laboratory, 1479 Gortner Ave, St Paul, MN 55108

^2^Department of Organismic and Evolutionary Biology, Harvard University, 26 Oxford St, Cambridge, MA 02138

^3^Herbarium and Center for Tree Science, The Morton Arboretum, Lisle, IL 60532

^4^Pritzker Laboratory for Molecular systematics and Evolution, The Field Museum, 1400 S Lake Shore Dr, Chicago IL 60605, USA

^5^School of Biological Sciences, University of Oklahoma, Norman, OK 73019

### Section S1: Figures

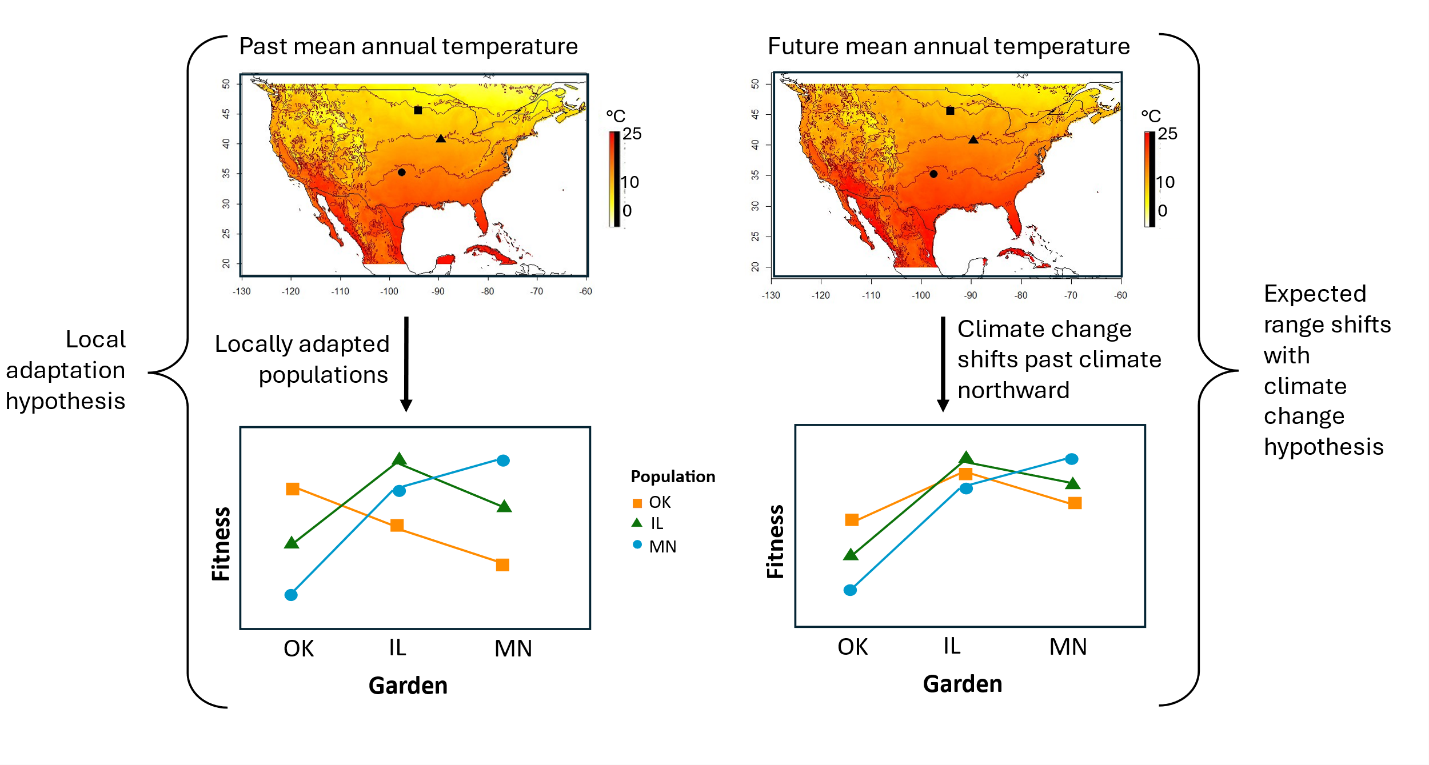

*Figure S1.* Illustrated hypotheses for local adaptation, adaptive differentiation, maladaptation, and shifting habitat suitability with climate change. The left panels represent classic local adaptation predictions where home populations have higher fitness than away populations with significant differences in fitness among populations in each garden, and each population has the highest fitness in its local garden. Populations have lower fitness in away gardens under the classic local adaptation hypothesis, indicating maladaptation. The right panels illustrate expectations under a warming climate, where populations still have higher fitness in their home garden compared to the away populations, but tend to have higher fitness in northern gardens compared to southern gardens due to shifting climate. These hypotheses do not take into account other environmental factors (including edaphic and microbial factors) that may be relevant to local adaptation.

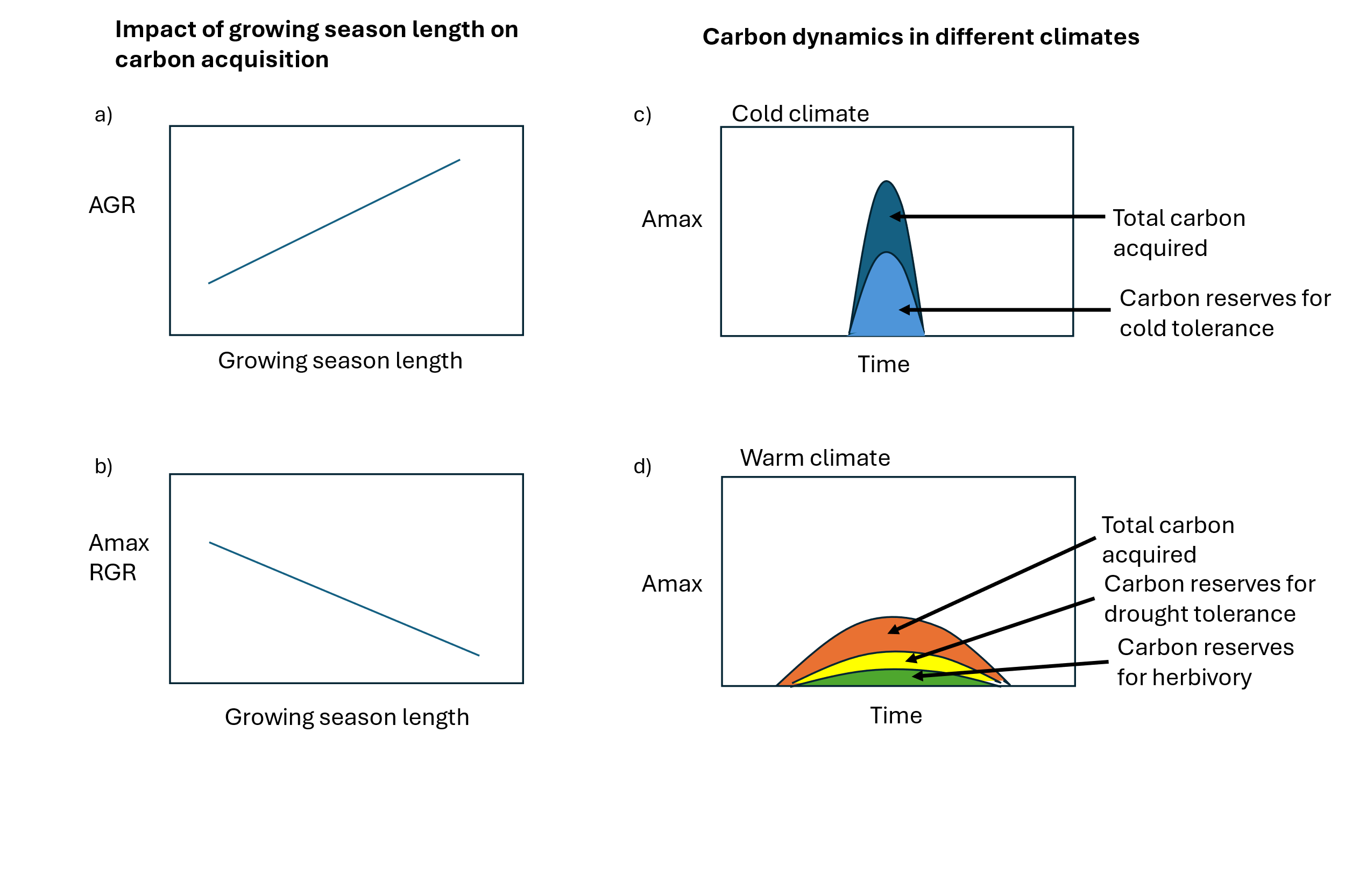

*Figure S2.* Predicted associations of growing season length with (a) absolute growth rate (AGR) and carbon assimilation rate (*A_max_*) and relative growth rate (b). Total growth over a growing season is expected to increase with a longer growing season, however, with longer growing seasons, the carbon assimilation rate and relative growth rate are expected to be slower and spread out throughout the longer growing season. We expect that carbon allocation will shift due to stressors introduced in colder (a) or warmer (b) climates. In cold climates, carbon acquisition is expected to occur at a fast rate over a short period of time allowed by the growing season. A portion of that carbon acquired is expected to be allocated toward non structural carbon reserves for cold tolerance. In warmer climates, carbon acquisition is expected to occur over a prolonged period over a slower rate. A portion of the carbon may be allocated toward drought tolerance or toward herbivore defense. The remaining carbon is available for allocation toward growth.

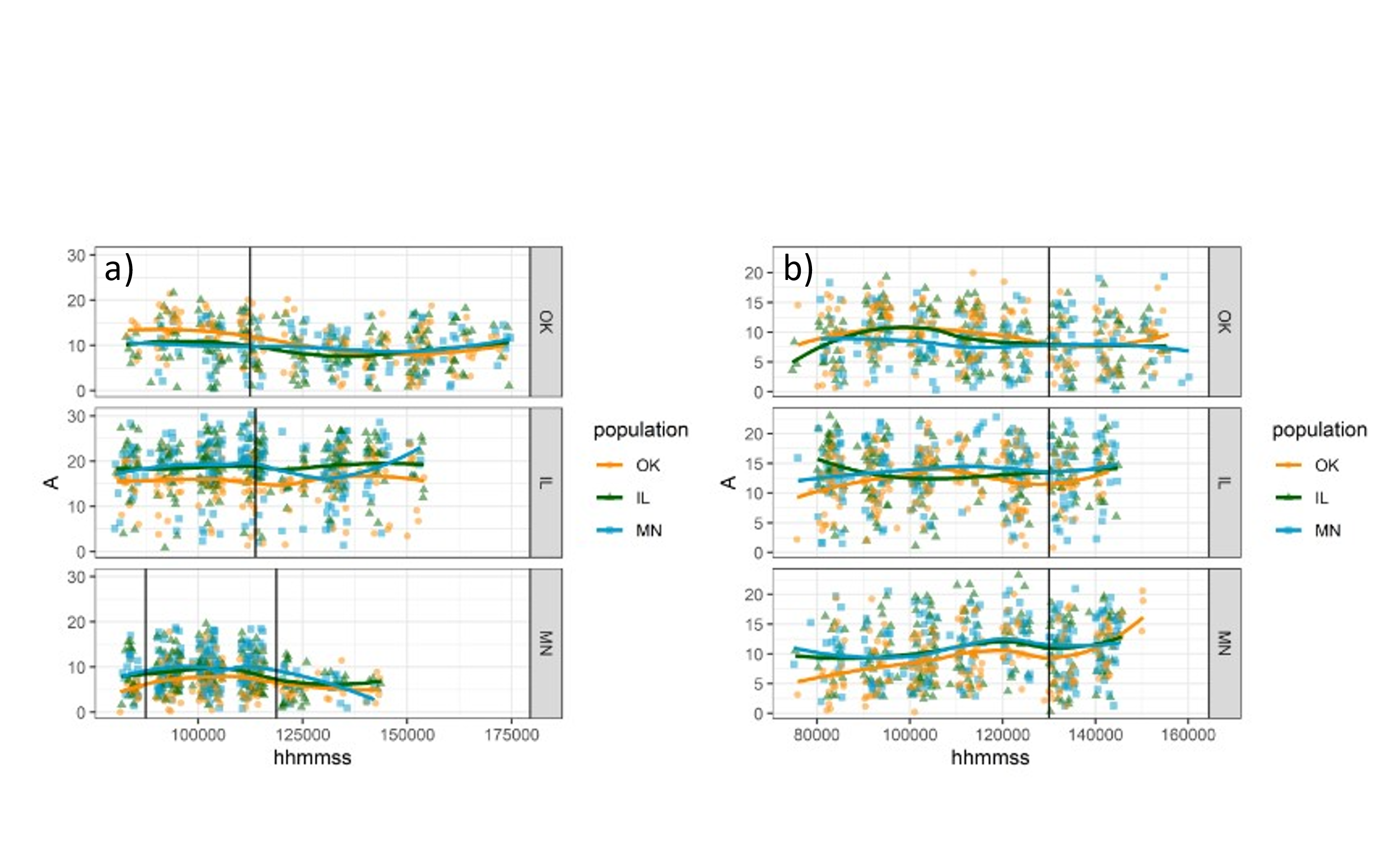

*Figure S3.* Diurnal curves for photosynthetic carbon assimilation (A) in each garden, with time of day on the x axis in 2023 (a) and 2022 (b). Each panel represents a garden, and each point represents an individual measurement. Black vertical lines indicate the cutoff points for A_max_. Populations are represented as orange circles for Oklahoma, green triangles for Illinois, and blue squares for MN. Colored lines indicate the trend in A for each population.

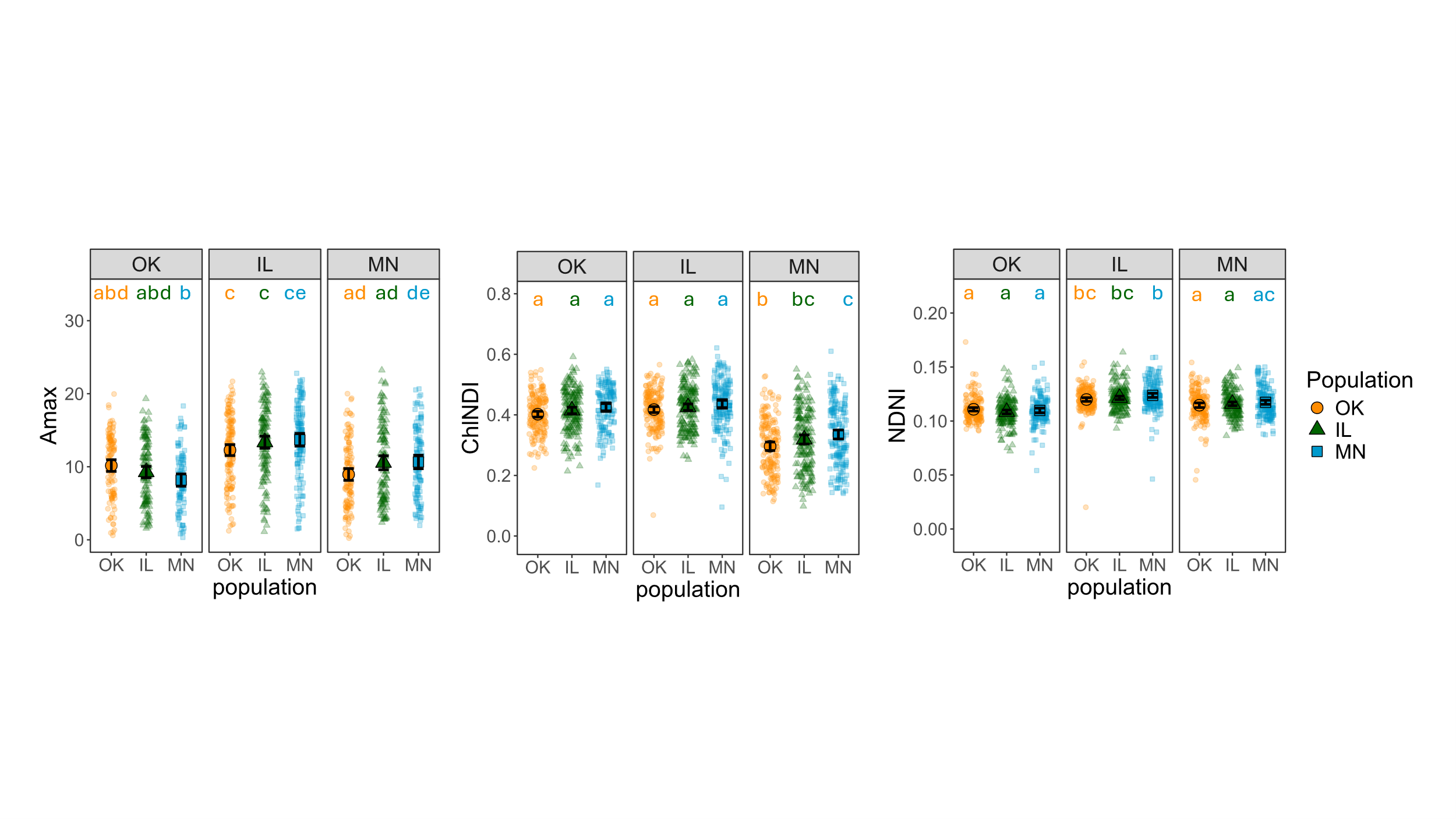

*Figure S4.* Physiological data from 2022 including the max photosynthetic rates (A_max_), chlorophyll normalized difference index (ChlNDI), and normalized difference nitrogen index (NDNI). Each panel represents a common garden located in Oklahoma (OK), Illinois (IL), or Minnesota (MN). The populations are represented by an orange circle for OK, a green triangle for IL, and a blue square for MN. Distinct letters denote statistical significance (p<0.05) for each population group.

### Section S2: Tables

*Table S.1.* Number of individuals for each measure for each population in each garden.

| **Measure** | **Garden** | **Population** | **Number of individuals** |
| --- | --- | --- | --- |
| A_max_ | OK | OK | 62 |
| A_max_ | OK | IL | 57 |
| A_max_ | OK | MN | 34 |
| A_max_ | IL | OK | 101 |
| A_max_ | IL | IL | 96 |
| A_max_ | IL | MN | 96 |
| A_max_ | MN | OK | 123 |
| A_max_ | MN | IL | 112 |
| A_max_ | MN | MN | 132 |
| ChlNDI | OK | OK | 61 |
| ChlNDI | OK | IL | 48 |
| ChlNDI | OK | MN | 42 |
| ChlNDI | IL | OK | 179 |
| ChlNDI | IL | IL | 183 |
| ChlNDI | IL | MN | 180 |
| ChlNDI | MN | OK | 157 |
| ChlNDI | MN | IL | 161 |
| ChlNDI | MN | MN | 165 |
| gswMAX | OK | OK | 62 |
| gswMAX | OK | IL | 57 |
| gswMAX | OK | MN | 34 |
| gswMAX | IL | OK | 101 |
| gswMAX | IL | IL | 96 |
| gswMAX | IL | MN | 96 |
| gswMAX | MN | OK | 123 |
| gswMAX | MN | IL | 112 |
| gswMAX | MN | MN | 132 |
| iWUE_max_ | OK | OK | 62 |
| iWUE_max_ | OK | IL | 57 |
| iWUE_max_ | OK | MN | 34 |
| iWUE_max_ | IL | OK | 101 |
| iWUE_max_ | IL | IL | 96 |
| iWUE_max_ | IL | MN | 96 |
| iWUE_max_ | MN | OK | 123 |
| iWUE_max_ | MN | IL | 112 |
| iWUE_max_ | MN | MN | 132 |
| NDNI | OK | OK | 61 |
| NDNI | OK | IL | 48 |
| NDNI | OK | MN | 42 |
| NDNI | IL | OK | 179 |
| NDNI | IL | IL | 183 |
| NDNI | IL | MN | 180 |
| NDNI | MN | OK | 157 |
| NDNI | MN | IL | 161 |
| NDNI | MN | MN | 165 |
| RGR stem volume | OK | OK | 167 |
| RGR stem volume | OK | IL | 140 |
| RGR stem volume | OK | MN | 103 |
| RGR stem volume | IL | OK | 185 |
| RGR stem volume | IL | IL | 189 |
| RGR stem volume | IL | MN | 194 |
| RGR stem volume | MN | OK | 184 |
| RGR stem volume | MN | IL | 188 |
| RGR stem volume | MN | MN | 182 |

*Table S.2.* Climate from 1940-2023 in each of the three gardens.

| **Garden** | **Mean annual temperature (°C) 1940-1960** | **Mean annual temperature (°C) 2003-2023** | **Δ mean annual temperature (°C) 1940-60 to 2003-23** | **Growing season temperature (°C) 1940-1960** | **Growing season temperature (°C) 2003-2023** | **Δ growing season temperature (°C)**  **1940-60 to 2003-23** |
| --- | --- | --- | --- | --- | --- | --- |
| Oklahoma | 15.6 | 16.75 | **1.14** | 26.78 | 27.71 | **0.93** |
| Illinois | 10.31 | 11.65 | **1.33** | 22.78 | 23.56 | **0.78** |
| Minnesota | 5.72 | 7.16 | **1.44** | 20.15 | 21.36 | **1.21** |

Table S.3. Average acorn volumes for each of the three populations*.*

| **Population** | **Average acorn volume (cm^3^)** |
| --- | --- |
| OK | 14.7 |
| IL | 3.5 |
| MN | 2.0 |

### Section S3: Methods supplement

**Acorn processing**

The acorns were processed by removing the caps and placing them in water, discarding acorns that floated as they were likely to be inviable. Viable acorns were placed in sand in cold storage until planting in the fall.

**Garden maintenance**

*Mulching*

In Illinois and Minnesota, the gardens were mulched at the end of the first growing season. In Oklahoma, mulching occurred in 2023 due to challenges sourcing mulching materials.

*Irrigation*

In Minnesota, for the first year after planting, the seedlings were watered five days a week for eight weeks, then two to three days a week for another four to five weeks. The second year, seedlings were irrigated once a week through July and August. Each time they were irrigated the seedlings were provided approximately 5 mm of water.

In Illinois, for the first two years of planting, the seedlings were watered only in weeks where < 1 inch of rain fell; supplemental watering aimed to provide 1 inch of watering in those weeks.

Similarly, in Oklahoma for the first two years of planting, the seedlings were watered twice a week for 30 minutes (approximately 1 inch per week total) any time there was not significant rain (>1 inch).

**Gas exchange parameters**

To keep the measurement conditions consistent across the three gardens, the chamber environment was maintained at 30 °C, with 60% relative humidity and 400 µmol/mol for the reference CO_2._ Light was set at PAR of 1200 µmol m^-2^ s^-1^, as was determined to be saturating based on light curves conducted in the Oklahoma garden on a subset of trees.

**Spectral data information and processing**

Our spectral measurements encompassed the wavelength range 340 to 2,500 nm with 1,024 spectral bands. The spectral sensors were matched by splicing at 990 nm and 1100 nm and interpolating 5 nm and 1 nm, respectively, using the match_sensors function in *spectrolab* package v. 0.0.18 (Meireles et al., 2021) in R v.4.3.0 (R Core Team, 2023).
